# Intrinsic neural excitability induces time-dependent overlap of memory engrams

**DOI:** 10.1101/2022.08.27.505441

**Authors:** Geoffroy Delamare, Douglas Feitosa Tomé, Claudia Clopath

## Abstract

Memories are thought to be stored in neural ensembles known as engrams that are specifically reactivated during memory recall. Recent studies have found that memory engrams of two events that happened close in time tend to overlap in the hippocampus and the amygdala, and these overlaps have been shown to support memory linking. It has been hypothesised that engram overlaps arise from the mechanisms that regulate memory allocation itself, involving neural excitability, but the exact process remains unclear. Indeed, most theoretical studies focus on synaptic plasticity and little is known about the role of intrinsic plasticity, which could be mediated by neural excitability and serve as a complementary mechanism for forming memory engrams. Here, we developed a rate-based recurrent neural network that includes both synaptic plasticity and neural excitability. We obtained structural and functional overlap of memory engrams for contexts that are presented close in time, consistent with experimental studies. Moreover, we showed that enhancing the initial excitability of a subset of neurons just before presenting a context biases the memory allocation to these neurons. We then explored the role of inhibition as a way of controlling competition among neurons from two ensembles. This work suggests mechanisms underlying the role of intrinsic excitability in memory allocation and linking, and yields predictions regarding the dynamics of memory engrams.

## Introduction

Neural circuits have the ability to form and retain memories that last from hours to years. In particular, pioneering anatomical studies (Scoville & Milner, 1957) have suggested that such circuits are located in the hippocampus, although they had long remained unobserved. Over the past decades, technological advances such as neural imaging and optogenetics allowed for the discovery of engram cells in multiple brain regions as the neural substrate for memory storage and retrieval (Josselyn & Tonegawa, 2020). They are defined as a subpopulation of neurons that is initially activated during presentation of a stimulus, followed by transient physical and/or chemical changes that lead to its specific reactivation during memory recall (Josselyn & Tonegawa, 2020). Engram cells have been observed in the hippocampus (Liu *et al*., 2012), in the amygdala (Rashid *et al*., 2016; Morrison *et al*., 2016) and the neocortex (Kitamura *et al*., 2017; Tonegawa *et al*., 2015). These studies have shed lights on the ability of neural populations to store and retrieve memories but the exact mechanisms responsible for the formation of memory engrams are not yet fully clear.

The mechanistic understanding of the formation and long-term stability of memory engrams has long been dominated by Hebbian learning (Hebb, 1949). Indeed, most computational models have focused on synaptic mechanisms, such as long-term potentiation (LTP) or depression (LTD) (Bliss & Collingridge, 1993; Josselyn & Tonegawa, 2020), that have been able to provide insight into the formation and stability of neural assemblies (Zenke *et al*., 2015; Litwin-Kumar & Doiron, 2014). As a result, the contribution of other important mechanisms, like intrinsic excitability (Titley *et al*., 2017), has remained underexplored. Indeed, previous experimental works have shown that neurons with high excitability are preferentially allocated to memory engrams (Silva *et al*., 2009; Han *et al*., 2007; Zhou *et al*., 2009). Interestingly, learning is known to transiently increase neural excitability, reducing the afterhyperpolarization of neurons over several hours (Thompson *et al*., 1996; Oh *et al*., 2003). This transient increase is likely due to the learning-induced expression of the transcription factor CREB (Rashid *et al*., 2016; Silva *et al*., 2009; Han *et al*., 2007), which is known to play a role in regulating neural excitability (Dong *et al*., 2006). As a result, time-varying excitability may account for overlapping neural ensembles encoding memories of events that are temporally linked (Sehgal *et al*., 2018), namely events spaced by a short temporal delay, as observed in the lateral amygdala (Rashid *et al*., 2016), the hippocampal dorsal CA1 (Cai *et al*., 2016; Shen *et al*., 2022), and the retrosplenial cortex (Sehgal *et al*., 2021). Previous theoretical works have described how the dynamics of plasticity-related proteins and excitability can lead to co-allocation of memories at the dendritic level (Kastellakis *et al*., 2016; Sehgal *et al*., 2021; Chowdhury *et al*., 2021). Attractor networks (Amit, 1989) have been previously used to describe the recurrent network properties of overlapping memory engrams (Gastaldi *et al*., 2021) but without taking excitability into account. Here, we developed a computational model that describes the formation of overlapping memory engrams, combining synaptic plasticity and activity-dependent intrinsic excitability. By focusing on the recurrent neural network dynamics and its link with behavior, we show that our model is able to explain - at the mechanistic level - experimental findings regarding overlapping neural ensembles and memory linking. Moreover, we uncover the potential mechanisms allowing neurons to compete for allocation to memory engrams, as observed experimentally. Our results suggest that the temporal linking of memory engrams arises from co-activation of different neural ensembles, mediated by the interaction of time-varying excitability and synaptic plasticity. Our model makes testable predictions about how the balance among inhibition, feed-forward inputs and excitability is crucial for determining the extent of overlap among engrams of temporally close events.

## Results

### Formation of a single memory engram in a recurrent network with excitability

In order to study the effect of excitability in memory allocation and linking, we built a rate-based model with feed-forward and recurrent connections, equipped with excitability and Hebbian plasticity. Excitability of each neuron *i* is modelled as a time-varying threshold *ε_i_* of the input-output function (Methods, Eq. 1). This excitability is initially sampled from a random distribution (Methods) and changes to a higher value when the neuron’s firing rate reaches a threshold *θ* before decaying to its initial value (Methods, Eq. 4). Feed-forward inputs are defined as a single layer divided in three subpopulations corresponding to different contexts. Feed-forward weights are set as a diagonal block structure to define three receptive fields (Methods, Fig. 1.a, Fig. S1), such that presenting a context increases the input current to a subpopulation of neurons in the main region. Recurrent connections are assumed to be all-to-all and plastic, according to a Hebbian learning rule (Methods, Eq. 2), and initialised at 0. We then stimulated the network by presenting different contexts (training phase, Methods).

**Figure 1:**
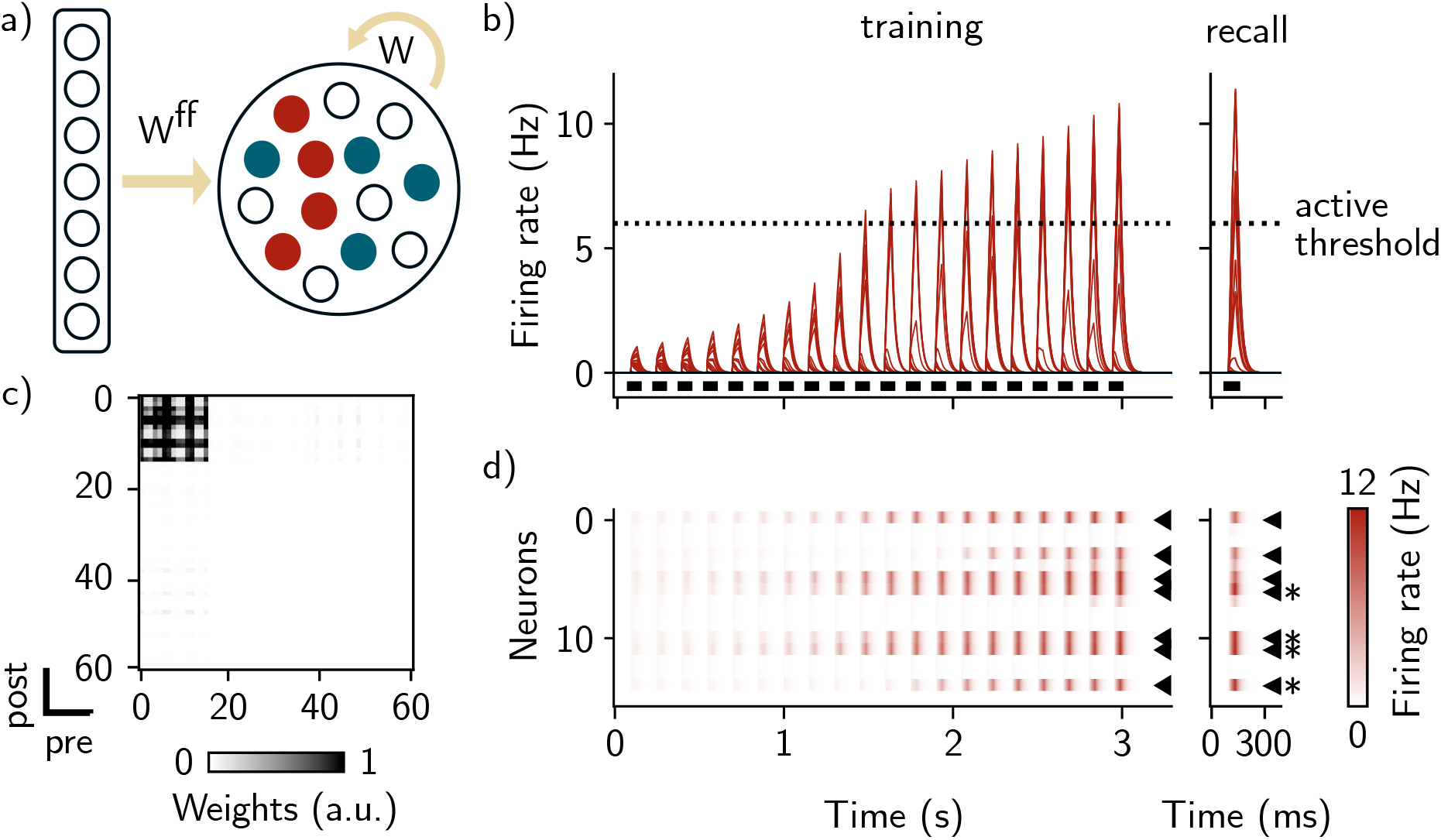
Encoding a single memory engram in a recurrent network equipped with intrinsic neural excitability. a) Diagram of the network architecture. Neurons in the input layer (left) project to the network with feed-forward connections *W*^FF^ (right). The feed-forward weights are defined such that neurons in the main layer have receptive fields (only two are shown here, red and blue neurons). b) Firing rates of all the neurons across time during training (left) and recall (right). Neurons responding preferentially to the context are shown in red while the other neurons are shown in black and do not respond to the stimulus. Black bars show presentations of the stimulus to the network. The dashed line is the “active” threshold, *i.e*. the threshold above which neurons are classified as active. c) Recurrent weights matrix after training. The block structure shows a neural assembly with stronger connections between neurons responding to the red context (0 to 15). d) Firing rate of each neuron responding to preferentially to the first context during training (left) and recall (right). Black arrows indicate 7 neurons that were tagged as “active” during training and that were reactivated during recall. During the recall phase, 4 of these 7 neurons (black stars) were stimulated.

We first observed that, after presenting the first context, the firing rates of neurons responding preferentially to this context (Fig. 1.b 0 to 3s) are above the “active” threshold θ, which we defined as the threshold above which neurons are classified as active. This was not the case for the other neurons in the network (Fig. 1.b, left). Analysing the recurrent weights matrix revealed that learning led to the formation of an assembly of neurons strongly connected to each other (Fig. 1c). The weights between neurons outside the assembly, however, have not significantly changed from their initial value equal to zero. We then sought to test the ability of the network to perform pattern completion. To this end, we stimulated the network with a partial cue and measured memory retrieval. We observed that stimulating 4 out of the 7 neurons composing the assembly, namely those that were tagged as active during training (Methods), is enough to activate all the neurons in the assembly (Fig. 1.d, right). This result shows that stimulating a subset of neurons of the assembly is sufficient to activate other neurons in the same assembly through strong intra-assembly connections.

### Intrinsic neural excitability induces overlap among memory engrams of temporally close events

Next, we investigated the effect of presenting a second context to the network either 6h or 24h after the first one (Fig. 2.a, Methods), inspired by previous experiments (Rashid *et al*., 2016). We designed our model in such a way that, after learning, excitability of neurons taking part in the newly formed assembly is increased, before slowly decaying to their baseline level (Methods, Eq. 4). Specifically, there is a transient increase in excitability after stimulating the network by the first context (Fig. 2.b, red triangle) and the second context (Fig. 2.b, blue triangle). We then measured memory recall for both contexts successively (Fig. 2.a). Presenting a second context after a 24h delay led to the formation of a second neural assembly in the recurrent weights matrix, distinct from the first one (Fig. 2.c, top, Fig. S2, a). This second assembly is composed of neurons that are responsive to only the second context. As in the previous section, neurons that are part of both assemblies are reactivated independently during memory recall (Fig. 2.d, top left and right). Interestingly, we found that if the second context is presented after a 6h delay, some off-diagonal weights are also reinforced for neurons responding preferentially to the second context (Fig. 2.c, bottom, Fig. S2, b). This suggests that the two memories are encoded by overlapping neural representations in the case where the contexts are presented 6h apart but not 24h apart. Indeed, when recalling the second memory, we observed a co-activation of neurons that take part in the first assembly in the case of a 6h delay (Fig. 2.d, bottom left and right). We can therefore quantify the overlap between the two assemblies, namely the number of neurons that were active during recall of both contexts, and found that it is higher in the case where the events were separated by 6h relative to 24h (Fig. 6, S6).

**Figure 2:**
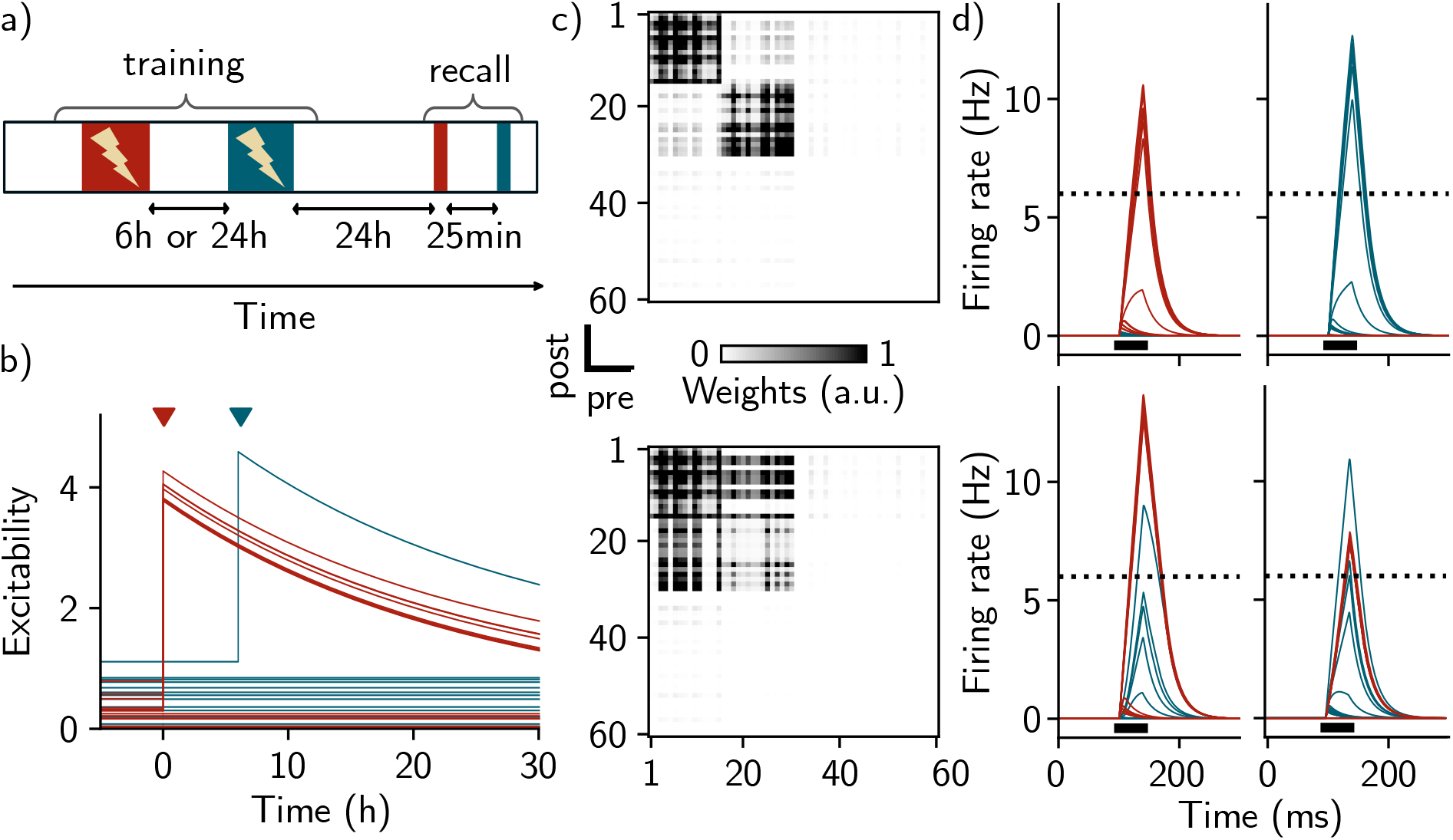
Training-induced increase in excitability induces overlap among memory engrams of temporally close events. a) Simulation protocol for studying the effect of forming two memories, spaced by different temporal delays. During the training phase, two contexts are presented 6h or 24h apart. After 24h, both contexts are recalled, separated by 25 min. b) Time course of excitability for each neuron of the network. Whenever the firing rate of a neuron crosses the active threshold, its excitability moves to a higher value (red and blue triangles corresponding to training on the first and second contexts, respectively), before decreasing to their initial value on a time scale of 24h (Methods). Red and blue traces correspond to neurons responding preferentially to the first and second contexts, respectively. c) Recurrent weights matrix immediately after training, in the case where the contexts are presented 24h apart (top) and 6h apart (bottom). d) Firing rates of individual neurons during recall of the first context (left) and the second context (right), in the case where the events are separated by 24h (top) and 6h (bottom). Neurons 1 to 15 respond preferentially to the first context (red) and neurons 16 to 30 respond preferentially to the second context (blue).

### Linking memories at the behavioral level in a fear conditioning simulation

We then asked whether this structural overlap among memory engrams could lead to memory linking. To that end, we modelled a fear conditioning experiment (Cai *et al*., 2016). We introduced freezing as an ideal-observer, namely a read-out value that is proportional to the sum of the neurons’ firing rate, integrated over the duration of the recall stimulus (Methods, Eq. 5). The unconditioned stimulus (US) was introduced as a multiplicative term in the Hebbian learning rule (Methods, Eq. 2) such that the recurrent weights are preferentially increased when the US is applied.

We presented three distinct contexts to the network, separated by 7 days and 5 hours (Fig. 3.a). In order to test memory linking, we presented the last context a second time, now paired with the US (Fig. 3.a, in blue) and measured the fear response in each of the three contexts. We observed that freezing was high when presenting either the blue context, which was paired with the US, or the yellow context, which was not paired with the US but was initially separated by 5h relative to the shocked context (Fig. 3.b). Conversely, presenting the red context, delayed by 7 days, elicited a freezing level comparable to the control case when no US was applied (Fig. 3.b). Our model was then able to show that two memories encoded close in time tend to be linked in such a way that recalling either memory can lead to a similar behavioral output, as shown experimentally (Cai *et al*., 2016; Shen *et al*., 2022).

**Figure 3:**
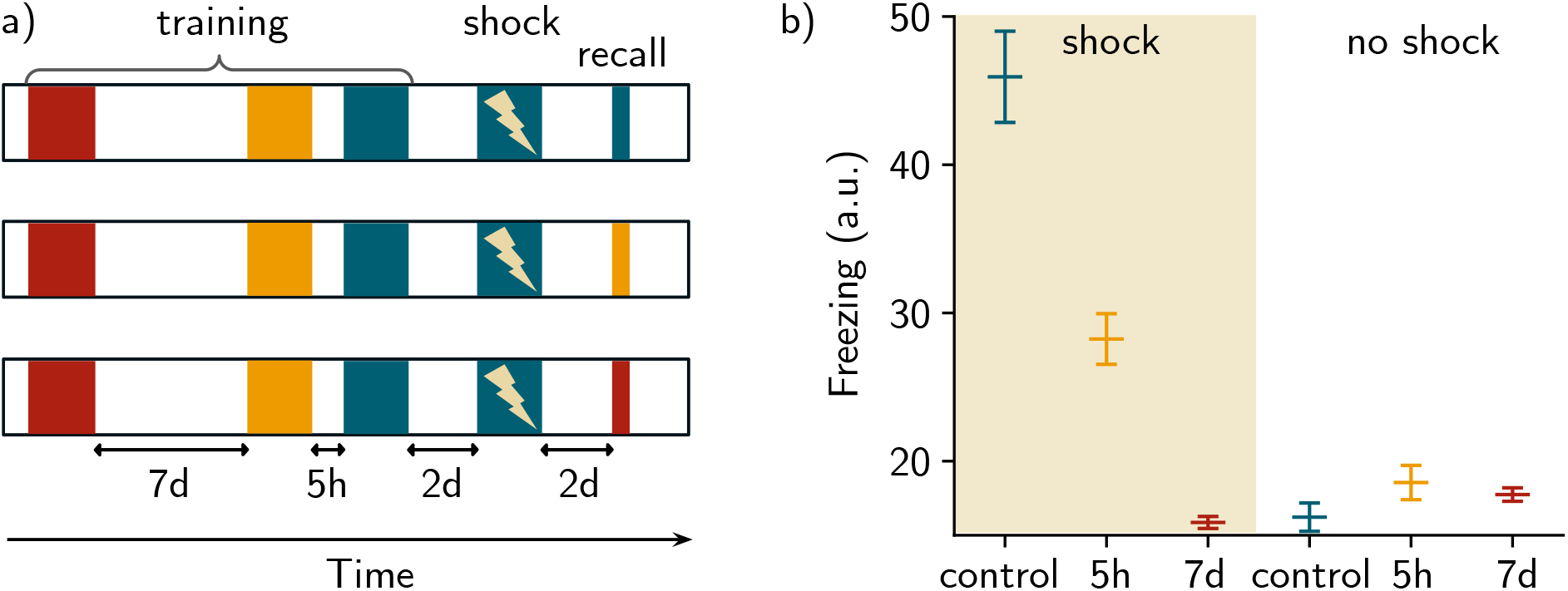
Memory linking in a fear conditioning simulation. a) Simulation protocol of fear conditioning. Three contexts are used and are delayed by either 7 days (red and yellow) or 5 hours (yellow and blue). Shock is then applied 2 days later in the blue context by pairing the context with an unconditioned stimulus (Methods). Memories are then recalled independently for each of the three contexts. b) Fear read-out upon recalling the three memories, when shock is applied (left) or not (right). n = 10 simulations and data are shown as mean ± s.e.m.

### Manipulating initial excitability biases neural allocation of memories

Given that excitability is a key mechanism for linking memory engrams, we then asked to what extent excitability could also play a role in biasing memory allocation. To that end, and inspired by previous experiments (Rashid *et al*., 2016), we increased the initial excitability of a subset of the neurons that respond preferentially to the first context (*ε*^+^, Fig. 4.a, Methods) during training of the first context. We then inhibited this subpopulation during memory recall (*i^+^*, Fig. 4.c, Methods) and measured the strength of memory recall (Freezing). We found that inhibiting the subpopulation whose excitability was enhanced reduced freezing during memory recall relative to the control case without manipulation of excitability (Fig. 4.d and Fig. S4.d). This suggests that neurons with enhanced excitability are preferentially allocated to memory engrams (Fig. S4.c, left).

**Figure 4:**
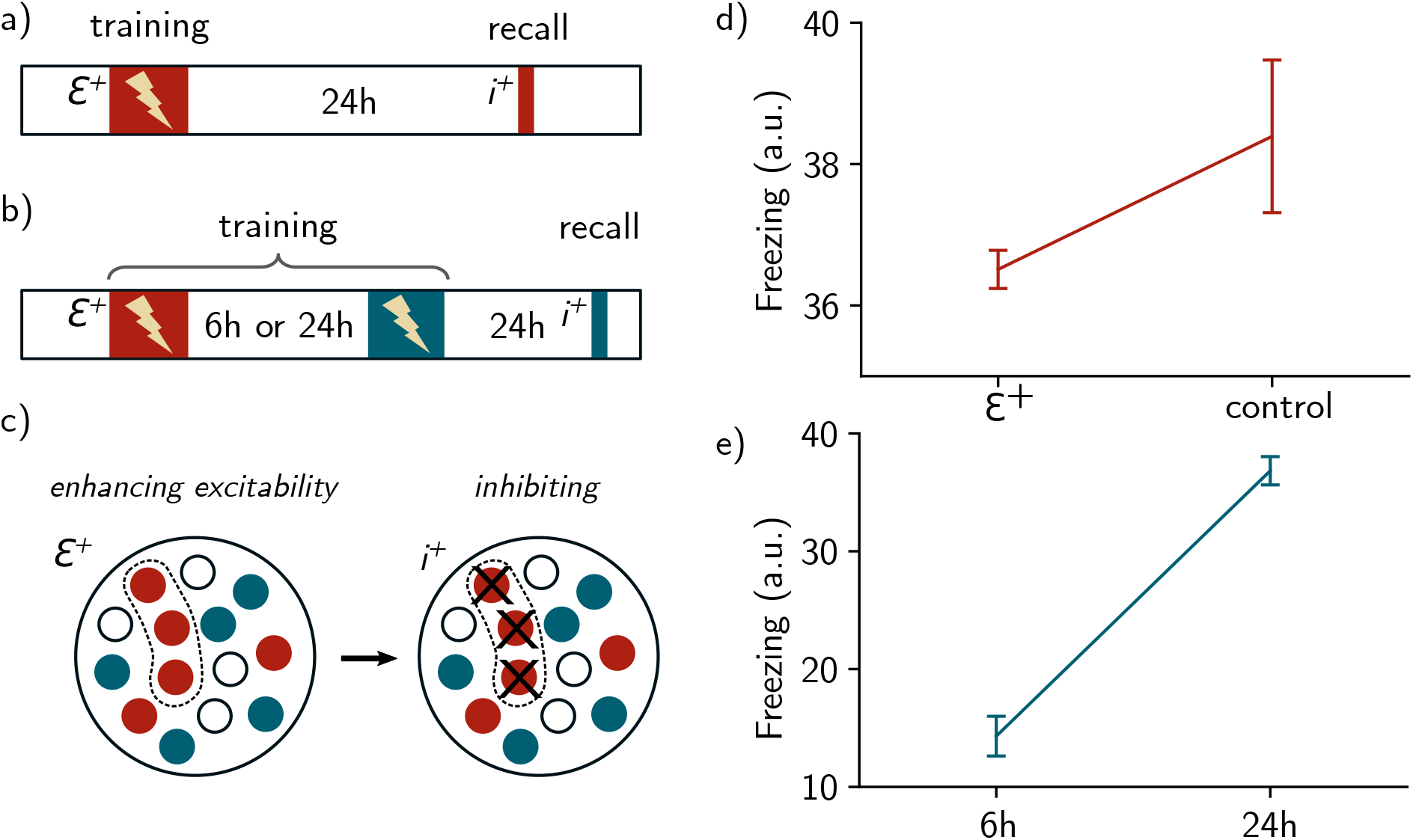
Neurons with enhanced excitability are preferentially allocated to memory engrams and overlapping ensembles. a) Protocol for biasing memory allocation to a subpopulation of neurons. Excitability of a subset of neurons is enhanced during training (*ε*^+^, top). During recall, this subpopulation is blocked (*i*^+^, Methods). In the control case, excitability is not manipulated. b) Protocol for biasing the overlap to a subset of neurons. Again, excitability is enhanced (*ε*^+^) for a subset of neurons during presentation of the first context (red). Then, a second context is presented (blue) after 6h or 24h, and the fear response to the second context is measured while blocking the subpopulation that received enhanced excitability (*i*^+^). c) Spatial representation of the protocol: a subset of *N_ε^+^_* neurons receives an enhanced excitability *ε*^+^, that is added to their initial excitability. During inhibition, *N_i^+^_* neurons from this subset are inhibited, receiving a negative current *i^+^* (Methods). d) Fear response to the context while blocking the subset of neurons, in the case where excitability is enhanced (*ε*^+^) or not (control). e) Fear response when recalling the second memory in b), when the two contexts are separated by either 6h or 24h. For all simulations, n = 50 simulations and data are shown as mean ± s.e.m.

Next, we presented two different contexts to the network,separated by 6h or 24h, and tested how increasing excitability to a subset of neurons during formation of the first memory could bias the overlap between the two memory engrams. We inhibited the subpopulation that received enhanced excitability during recall of the second context and we measured the fear response to the second context (Fig. 4.b and Fig. S4.b). In the case where the events were separated by 6h, inhibiting the subpopulation resulted in a reduction of freezing as compared to the case where the events were delayed by 24h (Fig. 4.e and Fig. S4.e). Given that the subpopulation *ε*^+^ is composed of neurons responding preferentially to the first context, this suggests that this subpopulation preferentially took part in the overlap between the two memory engrams (Fig. S4.c, right).

### Inhibition-induced competition among neurons crucially regulate memory allocation for temporally close events

Finally, we sought to evaluate how much neurons compete for memory allocation. To that end, we repeated the same protocol as in the previous section, but inhibiting the subpopulation (*i*^+^) during pre-sentation of the second context, instead of during recall (Fig. 5.a). We observed that the formation of the second memory was impaired when the contexts were presented 6h apart compared to the 24h delay (Fig. 5.b, solid lines). Note that this is the case whether or not the subpopulation (*ε*^+^) is inhibited (*i*^+^) during recall (Fig. S5.a-b, solid lines).

**Figure 5:**
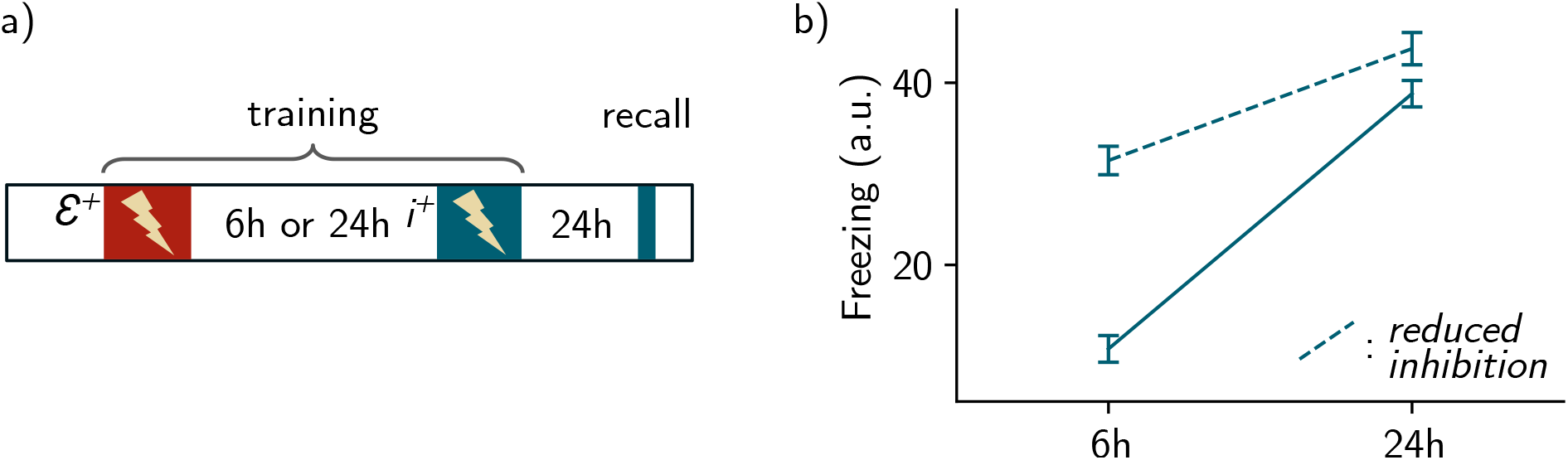
Neurons compete for allocation to memory engrams. a) Same protocol as Fig. 4.b but with the inhibition of the subset (*i*^+^) applied during presentation of the second context. b) Fear measurement when recalling the second memory when the events are separated by either 6h or 24h. The dashed line corresponds to the case where global inhibition *I*_1_ is reduced (Methods). For all simulations, n = 50 simulations and data are shown as mean ± s.e.m. (9 trials were excluded, Methods).

We then hypothesised that this memory impairment was driven by inhibition. We repeated the simulation as above while reducing the amount of inhibition in the network (Methods) during presentation of the second context, as inspired by previous experiments (Rashid *et al*., 2016). We observed that freezing was less impaired by inhibition of the subpopulation (*i^+^*) when the network inhibition was reduced (Fig. 5.b, dashed lines). Indeed, freezing in response to the second context increased more for a delay of 6h compared to 24h, relative to the case with baseline inhibition (Fig. 5.b and Fig. S5.b, dashed lines).

### The balance among inhibition, feed-forward inputs and excitability is crucial for forming overlaps

Overall, we found that excitability can induce overlap between memory engrams. This overlap is dependent on the temporal delay between the two contexts in a manner consistent with experimental findings in the lateral amygdala (Rashid *et al*., 2016) and in the hippocampal dorsal CA1 (Cai *et al*., 2016). Our result predict that the engram overlap arises from reactivation of the first ensemble when forming the second memory (Fig. S2.b). Indeed, co-activation of neurons encoding the first memory (red traces) along with neurons responding preferentially to the second context (blue traces) lead to strengthening of the weights between these two ensembles, due to Hebbian plasticity (Fig. 2.c, Fig. S2.b). We varied the temporal delay between the two contexts and found that the amount of overlap decreases when this delay increases.

We also predict that increasing the level of inhibition *I*_0_ leads to a decrease in the overlap between the two ensembles. Indeed, if the two events are separated by 6h, increasing *I*_0_ leads to a decrease in the overlap as compared to the control case (Fig. 6.c, 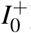). We found that decreasing the excitability decay timescale *τ_ε_* also leads to a decrease in the overlap (Fig. 6.c, 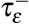). Indeed, the excitability increase following learning needs to be above a threshold otherwise the first ensemble cannot be reactivated even if the second context is presented after 6h (Fig. 6.e, Fig. S6.h). Finally, we also predict that the network can only form overlapping memory engrams if the feed-forward weights that do not form receptive fields 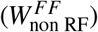 are within a defined range (Fig. S6.g). If these weights are too low, neurons that are not preferentially activated by the second context cannot be reactivated when presenting this context after 6h. On the other hand, if they are too high, the ensembles overlap independently of the temporal delay between the two contexts. In that case, we even observed an overlap with the novel context (Fig. S6.g, yellow line).

**Figure 6:**
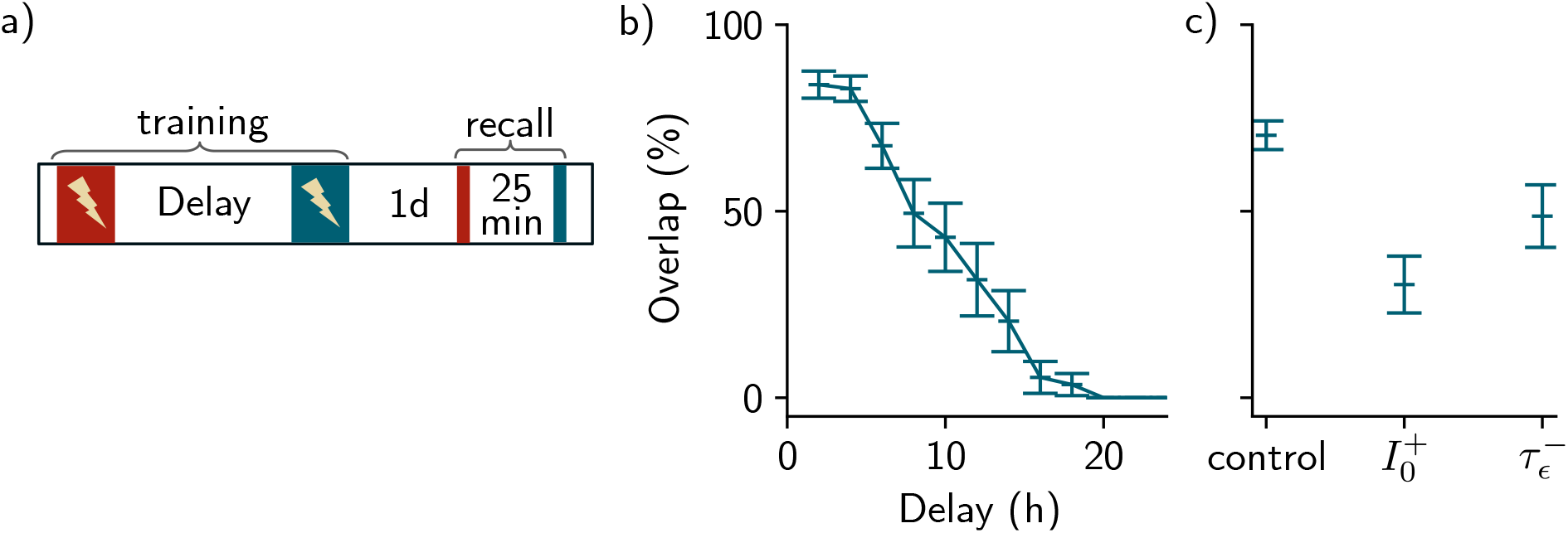
Predictions regarding the overlap among memory engrams. a) Protocol for forming overlap among memory engrams. During training, two contexts were presented, separated by a given temporal delay. The recall protocol allows for measuring the amount of overlap between engrams associated to the first context (red) and the second context (blue). b) Overlap among engrams against the temporal delay between the contexts. For each temporal delay, n = 20 simulations (3 were excluded, Methods). Results are shown as mean ± s.e.m. c) Overlap obtained for a 6h delay in the control case, the case where inhibition was increased 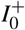 and the case where the excitability decay time was decreased 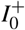. For the control case, n = 50 simulations (5 were excluded, Methods) and for 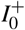 and 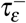, n = 20 simulations. Data are shown as mean ± s.e.m.

## Discussion

### Learning a single memory

We first showed that our network is able to form memories when stimulated by a feed-forward input. We attributed this formation to synaptic plasticity, independently from the dynamics of neural excitability. Indeed, the time scale of excitability is much slower than the time scale of Hebbian plasticity in our model, suggesting that the initial activation of the neurons by feed-forward inputs leads to strengthening of the synaptic weights through Hebbian learning. The structure of the neural assembly holding the formed memory is similar to previous theoretical works that have used attractor networks (Gastaldi *et al*., 2021; Amit, 1989). These neural assemblies are formed during the learning phase and are reactivated during memory recall. Importantly, memories can be recalled even when stimulated by a partial cue, suggesting that the neural activity is driven by recurrent connections. In particular, the structural change that leads to the formation of neural assemblies leads to the formation of a memory, as suggested in previous definitions of engram cells (Josselyn & Tonegawa, 2020).

The ability of the network to perform pattern completion is complemented by its ability to perform pattern separation. However, we did not evaluate pattern separation in our model. This would be interesting in particular because a recent study has shown that increased neural excitability in dendate gyrus improves pattern completion and separation (Pignatelli *et al*., 2019). The role of the recall-induced increase in excitability could explain this improvement. This could be directly tested in our model, for example by presenting a conflicting cue to the network and measuring pattern separation after recall of the memory.

Finally, our model does not take into account the evolution of memory engrams across time. Indeed, when the network is not stimulated, the recurrent weights are kept static. In this framework, the delay between the training phase and the recall phase has no impact on the recurrent weights.

### Overlap between memory engrams of temporally close events

Here, we built a model that is able to reproduce the overlap among memory engrams of events that are temporally linked. These overlaps have been observed in the amygdala (Rashid *et al*., 2016), the hippocampus (Cai *et al*., 2016) and the retrosplenial cortex (Sehgal *et al*., 2021) while it has been reported that these three regions are involved in a memory consolidation process known as systems consolidation (Tonegawa *et al*., 2018; Kitamura *et al*., 2017). However, it remains unclear what information is transferred from one region to another and investigating a potential transfer of overlap between brain regions would help understand how temporal memory linking evolves over the course of systems consolidation.

### Structural overlap of memory engrams induces memory linking

After showing that memories of temporally close events are structurally linked in overlapping neural ensembles, we showed that this overlapping structure can induce memory linking. In line with recent experimental studies (Cai *et al*., 2016; Yokose *et al*., 2017), we observed in our model that this overlap supports memory linking as the fear associated with one context can be transferred to another context (Fig. 3, Fig. S3).

We also note that this memory linking is a result of the short temporal delay between contexts. Indeed, in contrast to previous studies (Gastaldi *et al*., 2021; de Sousa *et al*., 2021), the overlap between two memory engrams is independent from the conceptual relation between the contexts in question, which we did not consider here. However, it is possible that this overlap supports the formation of mnemonic structures, as they have been observed in the hippocampus for instance (Barron *et al*., 2020; Deuker *et al*., 2016). Further work could be done to investigate the importance of overlapping memory engrams for more complex cognitive processes such as inferential reasoning (Barron *et al*., 2020; Zeithamova *et al*., 2012).

### Manipulating excitability biases neural allocation of memories

In our model, memory allocation is determined by two main factors. On the one hand, engrams are preferentially allocated to neurons that receive increased feed-forward inputs, as defined in the feedforward weights (Fig. 1). On the other hand, we showed that memory allocation is also biased towards neurons with high excitability (Fig. 4), consistent with previous studies (Zhou *et al*., 2009; Rashid *et al*., 2016). However, for the sake of simplicity, we did not explicitly probe the relative importance of the feed-forward weights compared to intrinsic excitability. To that end, it would be necessary to introduce some variability in the structure of the receptive fields, and subsequently investigate how neuronal memory allocation is impacted by feed-forward inputs versus excitability dynamics.

Here, we considered that excitability is mainly governed by the transcription factor CREB. We modelled its dynamics by increasing excitability instantaneously after learning and then allowing it to decay over a time scale of a few hours, as motivated by several experimental studies (Moyer *et al*., 1996; Oh *et al*., 2003; Thompson *et al*., 1996; Kitagawa *et al*., 2017). Although the results might be similar, it is important to note that these dynamics are conceptualized and that other mechanisms that are not considered here are also known to regulate neural excitability. For instance, internalisation of Kir2.1 channel increases neural excitability during memory recall (Pignatelli *et al*., 2019) while the expression of the C-C chemokine receptor type 5 (CCR5) is known to decrease excitability (Shen *et al*., 2022; Zhou *et al*., 2016). Adult-born neurons are also known to be more excitable than their counterparts (Silva *et al*., 2009). Finally, memory allocation may also be influenced by other mechanisms beyond the scope of the present study such as synaptic tagging and spine clustering (Rogerson *et al*., 2014; Kastellakis *et al*., 2016). We also showed that artificially increasing excitability in an ensemble of neurons could also bias co-allocation of this ensemble to further memories as shown in previous experimental findings (Rashid *et al*., 2016). This result arose naturally in our model because neurons with enhanced excitability are preferentially allocated to the first memory, and will then overlap with the second engram (Fig. S4.c).

### Role of inhibition in memory allocation and linking

Finally, we showed that neurons can compete for memory allocation and that the outcome of this competition is determined by both the initial excitability of neurons and the amount of inhibition in the network. We first showed that blocking neurons which received enhanced excitability during presentation of the first context impaired learning of a second context presented shortly after, suggesting that these neurons have a competitive advantage over the others for memory allocation. Secondly, we showed that reducing inhibition restored the ability of the network to learn the second memory, suggesting that competition is driven by inhibition.

In our model, neurons with a higher initial excitability are favoured for memory allocation and inhibit the remaining neurons, preventing them from taking part in a memory engram. This process has been previously shown experimentally (Rashid *et al*., 2016; Han *et al*., 2007) and this study provides a computational model that sheds light on the underlying competitive mechanism. Finally, we use a homogeneous global inhibition model, but further studies could explore the effect of populations of different inhibitory cell types on engram overlap and memory linking.

### Conclusion

In summary, we have built a recurrent neural network model that can reproduce the experimentally-observed neuronal overlap between temporally-linked memory engrams by combining both synaptic plasticity and neural excitability. Our results suggest that engram overlaps are crucially determined by the balance among inhibition, feed-forward inputs and excitability.

## Methods

### Rate model

Our rate-based model consists of a single recurrent neural network of *N* neurons (with firing rate *r_i_*, 1 ≤ *i* ≤ *N*) which receives inputs from an external region of *N*^in^ neurons (with firing rate 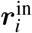, 1 ≤ *i* ≤ *N*^in^). The weights between the input region and the network are given by the matrix *W^FF^* (Fig. S1). Recurrent connections are given by the weight matrix W. Inhibition is introduced as 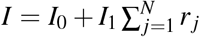, where *I*_0_ sets a baseline inhibition level and *I*_1_ scales an inhibition term proportional to the sum of the firing rates of the *N* neurons. Finally, excitability is added as a time-varying threshold *ε_i_*(*t*) of the input-output function. The rate dynamics of a neuron *i* is therefore given by:

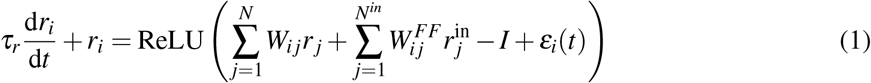

where *τ_r_* is the decay time of the rates and ReLU is the rectified linear activation function. In Fig. 5 and Fig. S5, the dashed lines correspond to the case where inhibition is reduced, *i.e. I*_1_ is set to a lower value 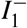 (Table of Parameters).

### Weights dynamics

The feed-forward weights *W*^FF^ are static and define three receptive fields (RF) that model three different contexts (Fig. S1). Neurons 1 to 15 respond preferentially to the first context, neurons 16 to 30 to the second context, and neurons 31 to 45 to the third context. All-to-all recurrent connections *W* are plastic and the weights *W_ij_* between each presynaptic neuron *i* and postsynaptic neuron *j* follow a Hebbian rule given by:

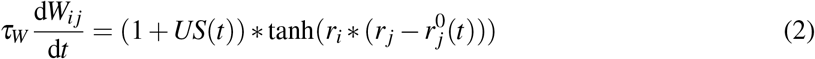

where *τ_W_* is the learning time constant and *US*(*t*) is the unconditioned stimulus (US) which is equal to *US*^+^ when US is applied (synchronously with stimulation of the context) and 0 otherwise. 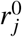 is the temporal mean over a time window *δ* of the firing rate of neuron *j*, given by:

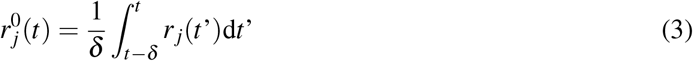

An upper cap *W*_max_ and a lower cap *W*_min_ are applied to the recurrent weights *W* to prevent them from being negative or too high.

### Intrinsic neural excitability

Intrinsic neural excitability follows dynamics that have been previously hypothesised to be due to the increase in the CREB transcription factor following learning (Silva *et al*., 2009). Each neuron’s initial excitability 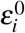 is sampled from a half-normal distribution of mean 0 and standard deviation 0.5. If the firing rate of a given neuron *i* reaches a set active threshold *θ*, its excitability *ε_i_* moves from its initial value 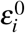 to a higher value *E* before decaying to 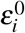 with a time scale *τ_ε_*:

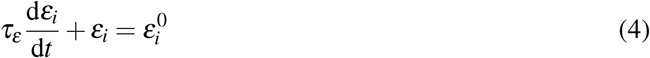

Note that we did not consider any increase in excitability following recall.

### Stimulation protocol

During training, either the first, second or last third of the *N*^in^ neurons from the input region are activated, namely their firing rates are set to a fixed value r_CS_. This activation is repeated *N*_stim_ times for a duration Δ*T*, with an interstimulus delay Δ*S*. During the recall period, one of the three contexts is presented for a duration Δ*T*. During training in all figures (except Fig. 3 and Fig. S3) shock is applied synchronously with the context, namely the value *US* in the learning rule (Eq. 2) is set to a non-zero value *US^+^*. In Fig. 3 and Fig. S3, shock is applied during the last presentation of the blue context (when specified).

During the simulations where excitability is manipulated (Fig. 4, 5, Fig. S4 and Fig. S5), the first *N_ε^+^_* neurons received an enhanced excitability *ε^increase^*, that is added to their initial excitability *ε*^0^ during presentation of the first context. Then, when inhibition is applied (*i*^+^), the first *N_i^+^_* neurons receive an external negative current *i*.

### Behavioral read-out

We introduced a read-out variable in order to compare it with the freezing levels measured in experiments. To that end, we modeled freezing using an ideal observer defined as:

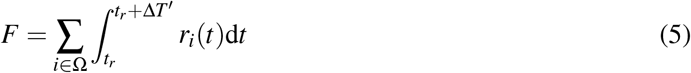

where *t_r_* is the onset of the recall stimulation and Δ*T’* = 100 ms is the integration time, corresponding to the temporal window during which the neuron is active. Ω denotes the ensemble of neurons belonging to the engram, namely the set of neurons crossing the active threshold *θ* during the recall time window.

The engram overlap (Fig. 6 and S6) is computed as the number of neurons responding to both recall of the first context and another context (either the same context, a novel context or a context that was presented with a 6h or 24h delay) divided by the number of neurons responding to the first context.

### Exclusion criteria

During some simulations, the firing rates of some neurons increased and reached non-realistic values. We defined a threshold of 100 Hz and decided to exclude any trials where the firing rate of any neuron reached 100 Hz at any time point. Around 10% of the trials were typically excluded.

### Integration

Integration was done using Euler’s method on Python. A time step of 0.5 ms was used during and 3 s after training sessions, and during and 300 ms after recall sessions. Between training and recall, a time step of 20 s was used.

### Table of parameters

An initial set of parameters was used in most of the figures except in Fig. 3 and Fig. S3. This initial configuration was chosen to match previous experimental results in the amygdala (Rashid *et al*., 2016). In Fig. 3, a second set of parameters was used to match engrams overlap measures observed in the dorsal CA1 (Cai *et al*., 2016).

| Parameter | Description | Fig. 1, 2, 4, 5, S1, S2, S3, S5, S6 | Fig. 3, S1, S4 |
| --- | --- | --- | --- |
| $N$ | Number of neurons in the main region | 60 | |
| $N^{\text{in}}$ | Number of neurons in the input layer | 30 | |
| $\tau_e$ | Excitability decay time constant | 24 h | |
| $E$ | Excitability jump | 3.5 | 4 |
| $\tau_r$ | Rate decay time constant | 15 ms | |
| $\theta$ | Active threshold | 6 Hz | 4 Hz |
| $I_0$ | Inhibition baseline | 6 | |
| $I_1$ | Inhibition multiplicative factor | 0.9 | |
| $I_1^-$ | Reduced inhibition multiplicative factor | 0.88 | |
| $\Delta T$ | Stimulation duration | 40 ms | |
| $N_{\text{stim}}$ | Number of stimulation per training sessions | 20 | 15 |
| $\Delta S$ | Interstimulus delay | 150 ms | |
| $W_{\text{RF}}^{\text{FF}}$ | Feed-forward weights corresponding to the RF neurons | 0.3 | |
| $W_{\text{non RF}}^{\text{FF}}$ | Feed-forward weights corresponding to the non RF neurons | 0.2 | |
| $\tau_w$ | Hebbian rule time constant | 750 ms | 500 ms |
| $W_{\text{max}}$ | Upper cap for plastic recurrent weights | 1 | |
| $W_{\text{min}}$ | Lower cap for plastic recurrent weights | 0 | |
| $US^+$ | Unconditioned stimulus strength | 1 | 0.5 |
| $r_{\text{CS}}$ | Firing rate of the neurons from the input layer when activated | 4 Hz | |
| $\delta$ | Averaging time of firing rates | 15 s | |
| $N_{\text{E}^+}$ | Number of neurons that receive enhanced excitability | 8 | - |
| $\epsilon^{\text{increased}}$ | Increase in excitability | 5 | - |
| $N_{\text{i}^+}$ | Number of neurons that are inhibited | 6 | - |
| $i$ | Inhibition strength | 0.5 | - |

## Acknowledgment

We thank Sadra Sadeh and Inês Completo Guerreiro for helpful comments on the manuscript, Yosif Zaki and Denise J. Cai for useful feedback and members of the Clopath lab for discussion and support.

## Supplementary information

**Figure S1:**
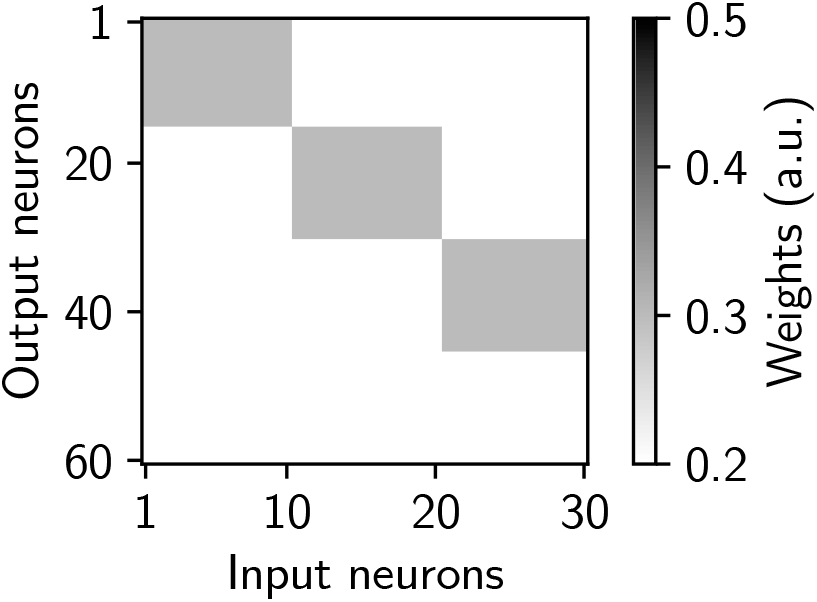
Feed-forward weights matrix. Three receptive fields are defined by strong weights in a block-diagonal structure. When presenting the first context for example, the first 10 neurons of the input region are activated which in turn stimulate preferentially the first 15 neurons in the main region.

**Figure S2:**
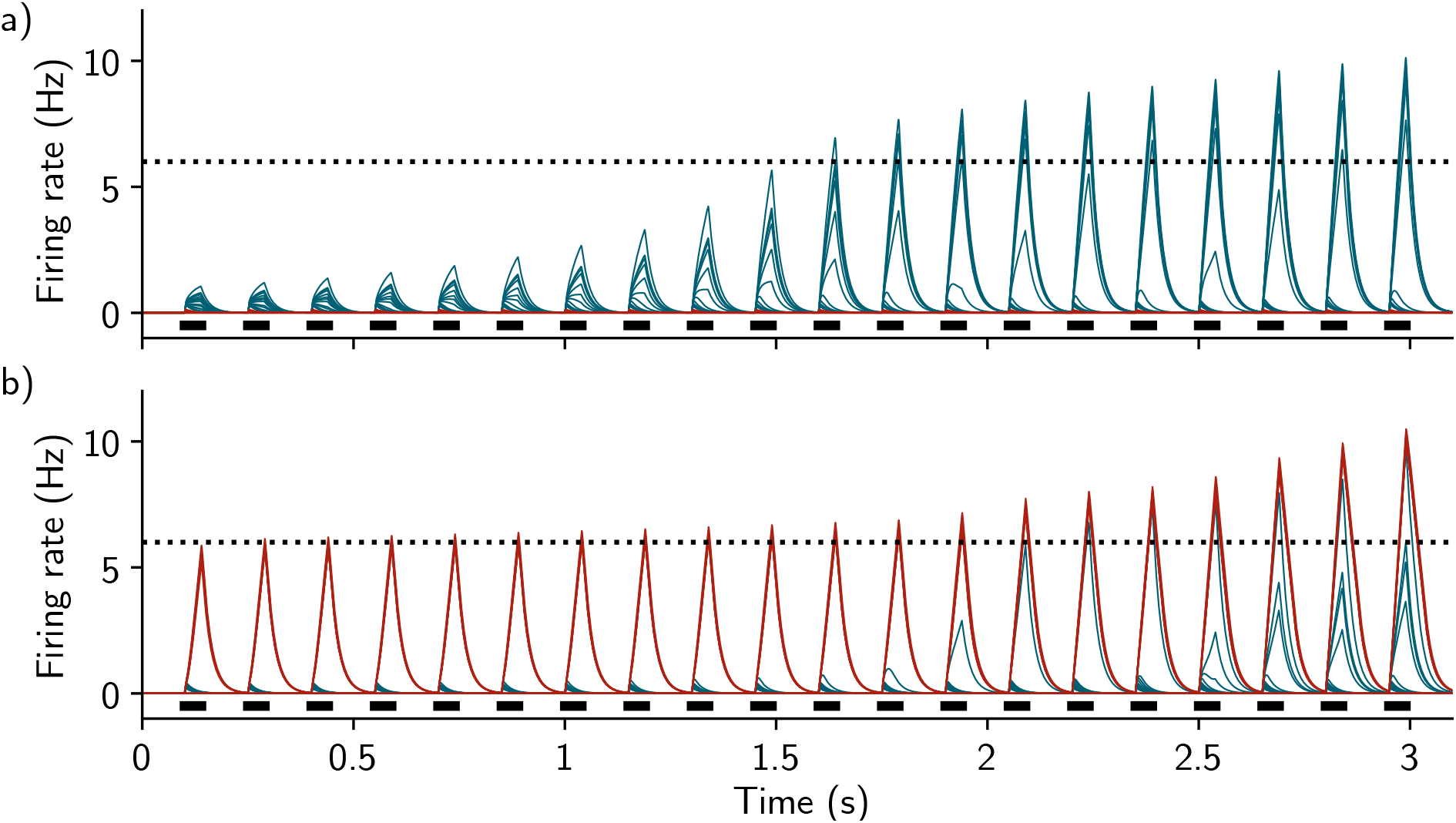
Firing rates of all the neurons as a function of time during presentation of the second context. a) Time course of the firing rate of all the neurons during presentation of the second context, when presented 24h after the first context (same protocol as Fig. 2). b) Same as a) when the second context is presented 6h after the first one. Blue and red traces correspond to neurons responding preferentially to the first and second context, respectively. The dashed line corresponds to the active threshold and the black bars to the stimulation.

**Figure S3:**
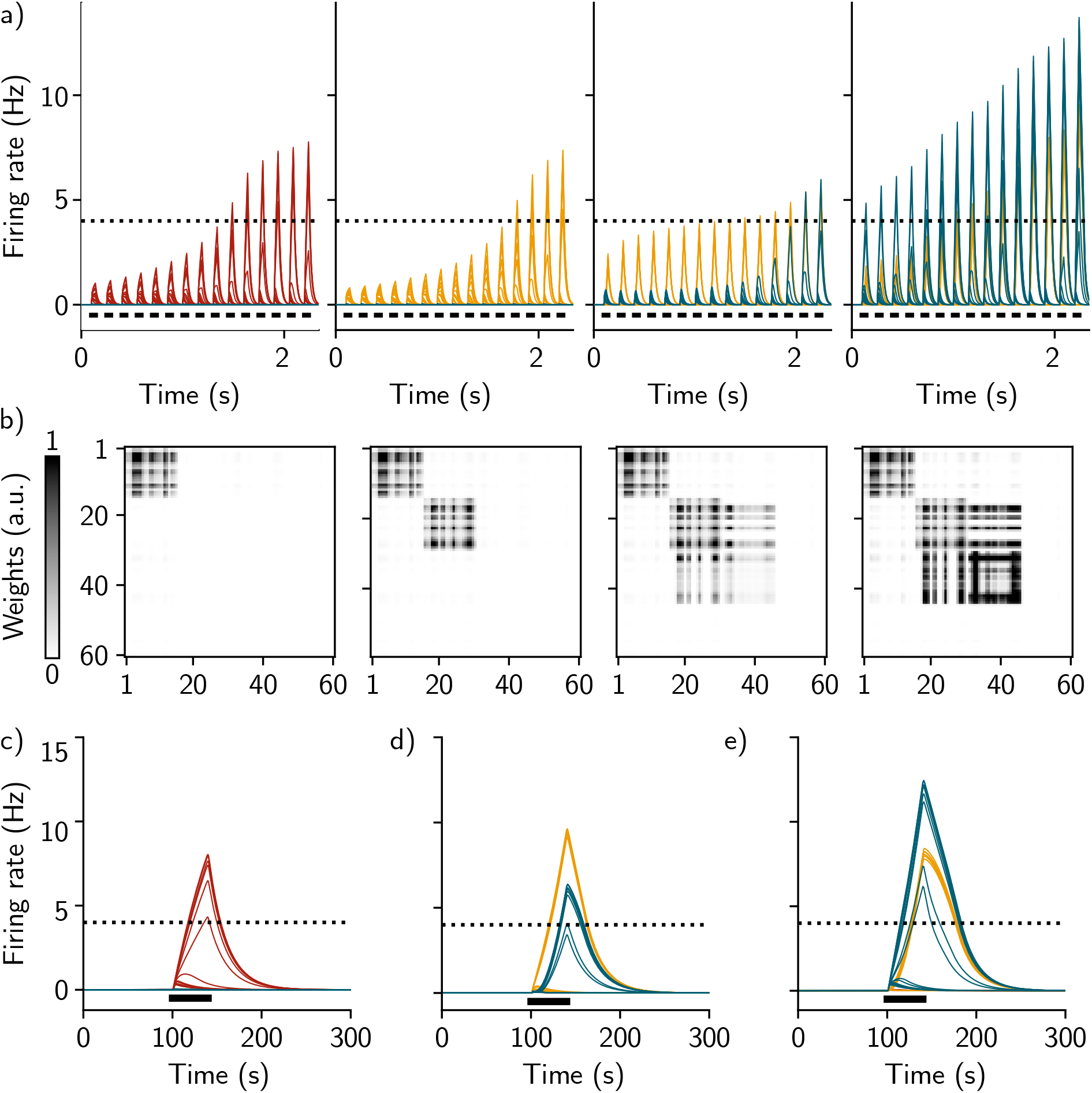
Network dynamics leading to memory linking. a) Firing rates of all the neurons across time during training in the fear conditioning protocol depicted in Fig. 3a. Each color represents a population of neurons that receives high input current when one of the contexts is presented. The first, second and third panels correspond, respectively, to the presentation of the first, second and third contexts, with a delay of 7 days (between the first and the second one) and 5 hours (between the second and the third one). Shock is applied during the second presentation of the third context, 2 days after the presentation of the third context (rightmost panel). The dashed line corresponds to the active threshold and the black bars to the stimulation. b) Corresponding recurrent weights matrices after presentation of each of the three contexts (first three panels) and after the second presentation of the third context paired with the shock (rightmost panel). c), d) and e) Firing rate of the neurons during recall of the first, second and third memory, respectively, 2 days after the shock. The dashed line corresponds to the active threshold and the black bars to the stimulation.

**Figure S4:**
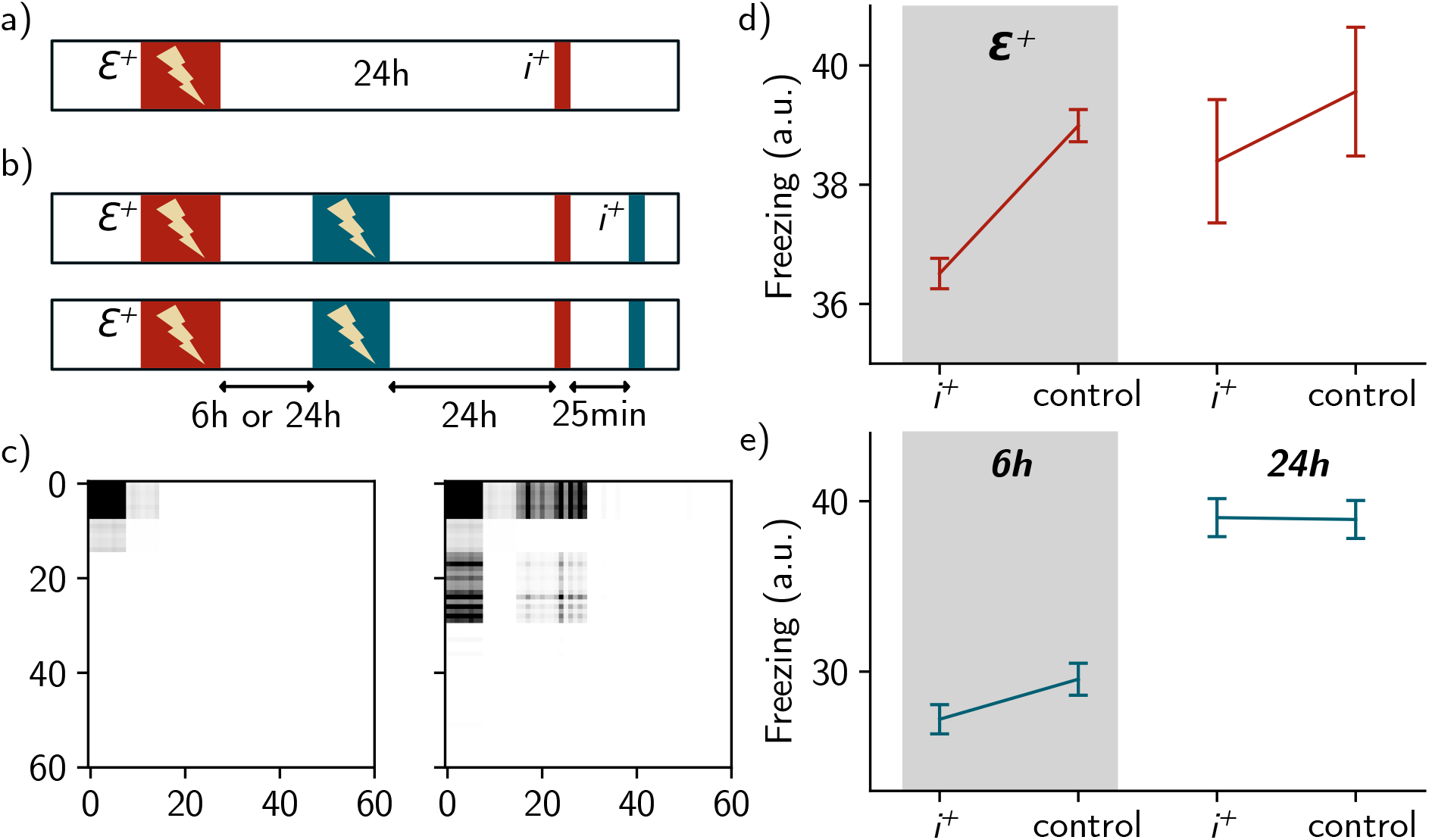
Extended manipulations to bias memory engrams allocation. a-b) Same as Fig. 4a and Fig. 4b where a subset of neurons receives enhanced excitability (*ε*^+^) or not and is either inhibited (*i*^+^) or not (control) during recall. c) Recurrent weight matrix after presentation of the first context (left) and the second context (right) for a 6h delay. d-e) Same as Fig. 4d and Fig. 4e, when the subset is either inhibited (*i*^+^) or not (control) during recall. For each conditions, n = 50 simulations and data are shown as mean ± s.e.m.

**Figure S5:**
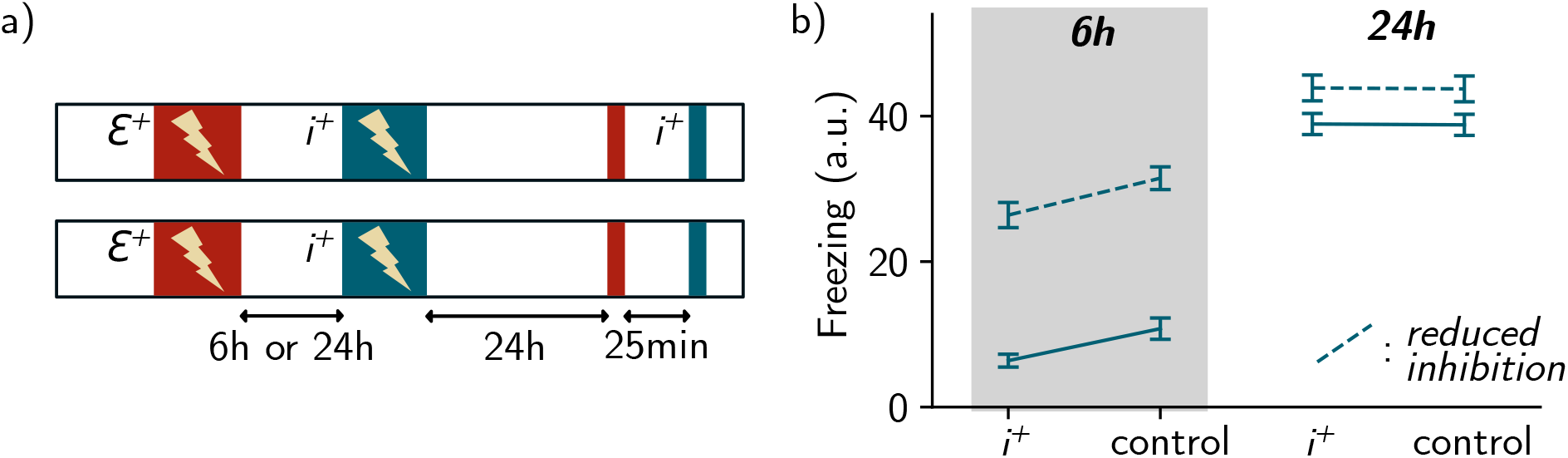
Extended manipulations to highlight competitive mechanisms. a) Same as Fig. 5a, when the subset either inhibited (*i*^+^) or not (control) during recall. b) Fear read-out upon presentation of the second context. Global inhibition *I*_1_ is either at baseline (*I*_1_ = 0.9, solid line) or reduced (*I*_1_ = 0.88, dashed line) during presentation of the second context. For each conditions, n = 50 simulations (9 were excluded, Methods) and data are shown as mean ± s.e.m.

**Figure S6:**
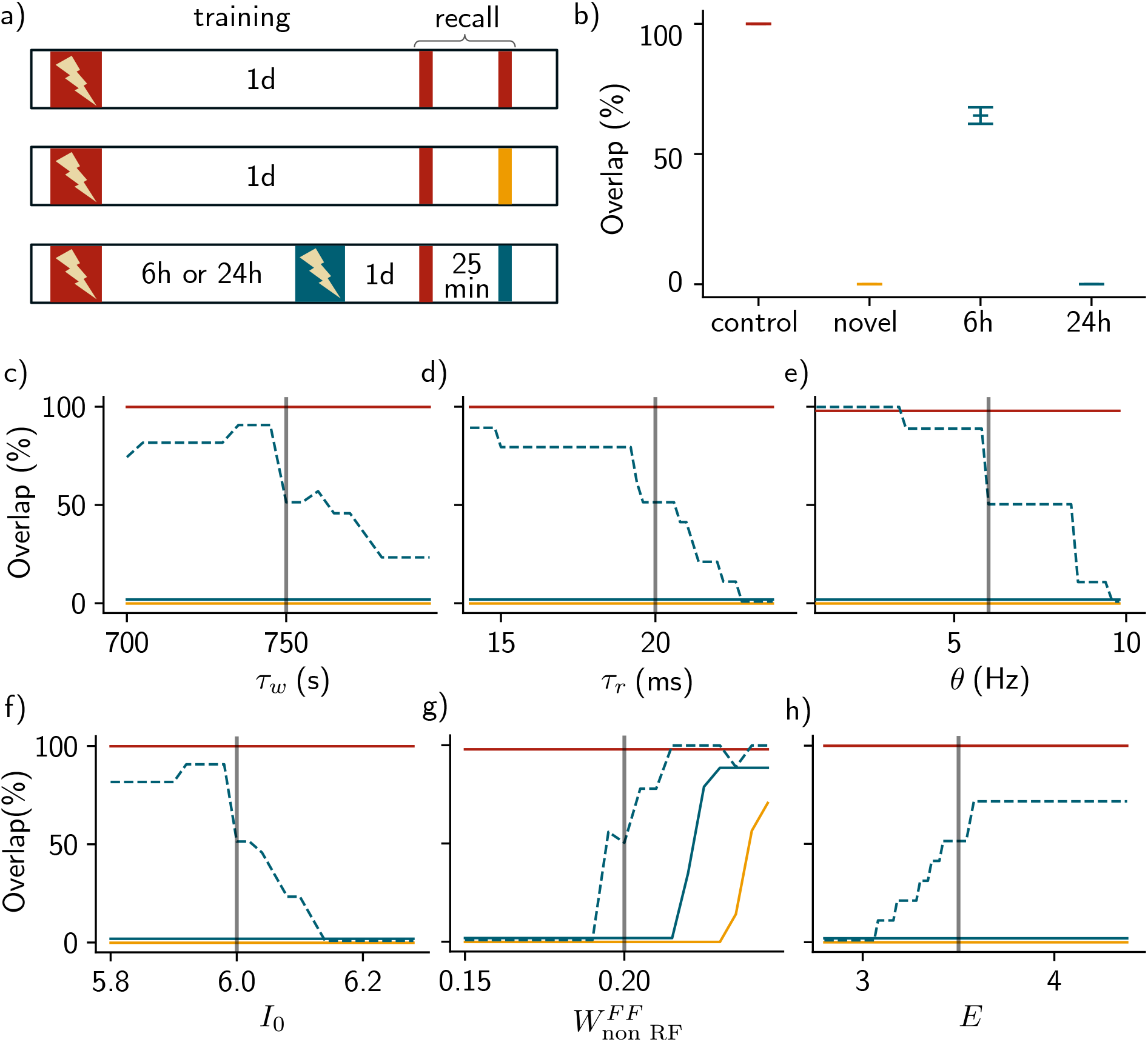
Model sensitivity analysis. a) Protocol for forming overlap among memory engrams. After training, the recall protocol allows for measuring the amount of overlap between engrams of the first context (red) and either the same context (top row, red), a novel context (middle row, yellow) or a second context (bottom row, blue), presented 6h or 24h after the first one. b) Overlap among engrams for the four cases in a): control case in red, novel context in yellow, 6h and 24h delays in blue. Here, the first set of parameters has been used (Methods). n = 50 simulations (5 were excluded, Methods) and data are shown as mean ± s.e.m. c-h) Overlap in each of the four cases in a) as a function of the main parameters of the model (Methods, Table of parameters). The blue dashed line corresponds to the 6h delay and the blue solid line to the 24h delay. Each grey line corresponds to the parameter that has been selected for the simulations in the main figures.

## Notes

### Competing Interest Statement

The authors have declared no competing interest.

